# Astrocytic Sleep Homeostasis Model

**DOI:** 10.1101/2022.10.23.513378

**Authors:** Ghanendra Singh

## Abstract

Sleep awake cycle is critical for cognitive and functional abilities. Conventional sleep homeostasis mechanisms are neuronal in nature and recent views indicate glial regulation of the sleep-wake process. Mechanisms of homeostatic regulation of sleep remain to be understood. A simplified astrocytic sleep-awake homeostasis mathematical model with sleep pressure and synaptic strength dynamics is proposed using feedback control loops. The model provides insights into the emergence of two discrete states through sleep and awake promoting neuronal populations giving rise to a homeostatic process S and oscillatory process C is regulated by astrocytic sleep pressure. It also explains the variations seen in synaptic strength dynamics during sleep and awake states.

## Introduction

Sleep plays an important role in cognitive functions of the brain, like learning and memory. Sleep deprivation leads to impaired cognition and sleep disorders. To understand the dynamics of sleep, initially a lotka-volterra based model proposed for reciprocal interaction between neuronal populations providing the basis of sleep cycle oscillations [1] which replicates the dynamics of sleep cycling over time. Stable oscillating solution and the limit cycle properties explained the observations under different conditions. Still, it did not explained in depth about the emergence of state transitions between sleep and awake states.

A two-process model [2] was proposed to understand the dynamics between sleep and awake states. It consists of a homeostat (Process S) describing sleep pressure interacting with a circadian pacemaker (Process C) to regulate sleep. Sleep pressure accumulates during wakefulness and dissipates during sleep following a circadian cycle [2]. Neuronal control of sleep regulation by measuring the sleep propensity and circadian rhythm sets the threshold for falling asleep or waking up [3]. Model provided a conceptual framework for qualitatively interpreting different experimental results in sleep research such as sleep deprivation, dysfunction and sleep disorders. However, it had its own limitations as a phenomenological model, such as in determining the thresholds for the sleep process for transitioning between states. How the process S and process C are related with each other? Nor it explains the different sleep states.

In sleep wake flip-flop model [4], neuronal populations play an important role in waking and sleep states. The waking state is maintained by wake-promoting neuronal population activity on the ascending arousal pathway [5]. Sleep-active neurons maintain the sleep state in the ventrolateral preoptic nucleus. Mutual inhibition between sleep and wake-active populations produces a state similar to a flip-flop electrical switch to maintain segregation and provide stability between the states. Flip-flop model provides a physiological substrate for sleep-wake transition mechanisms. Dynamic transition explains transition rates, stability of states, rapid transitions and loss of inhibition strength in sleep disorders. Though, it does not explain the condition required for the dynamic transition by which mutual inhibition between neuronal populations occurs. Also, intermediate states do exist in the brain disorders [6]. Both fast and slow transitions occurs in a continuous manner which cannot be explained by a simple binary switch and requires further revision.

Several other conceptual models have been proposed to account for different aspects of sleep regulation [7]. In sleep switch model, wake and sleep-promoting neurons inhibit each other, which results in stable wakefulness and sleep states. Disruption of wake or sleep-promoting states results in instability [4]. Previously published model accounts for several features of the human sleep-wake cycle [8]. The model demonstrates how these features depend on interactions between a circadian pacemaker and a sleep homeostat and provides a biological basis for the two-process model for sleep regulation. Another mathematical model for sleep-wake dynamics is the mutual inhibition model with mutual inhibition between sleep and awake promoting neuronal population regulated by homeostatic and circadian processes compared to two process models using two oscillators [9]. A framework to quantitatively model neural circuits and the role of feedback in sleep and arousal states was discussed by [10]. An integrative model for sleep/wake regulation in which an integrator neuron classifies the state as sleep or wake is hypothesized as an alternative to the flip flop model [11].

Sleep plays a regulatory role in synaptic plasticity [12]. Sleep synaptic hypothesis [13] states that wakefulness is associated with synaptic potentiation in several cortical circuits, Synaptic potentiation is tied to the homeostatic regulation of slow-wave activity. Slow-wave activity is associated with synaptic downscaling. Synaptic downscaling is tied to the beneficial effects of sleep on performance. A sleep regulatory subtance (SRS) TNF, is involved in synaptic scaling. The implications for sleep are that sleep-regulatory mechanisms cannot be separated from a sleep-connectivity stabilization function [14]. There are central global coordinators of sleep-wake states, such as the clock mechanisms of the suprachiasmatic nuclei. [15]. Sleep is regulated in a neuronal-assembly use-dependent manner. The mechanisms that underlie neuronal-assembly sleep include an enhanced activity of sleep-regulatory substances induced by neuronal use-enhanced metabolism. [16]

A mathematical model of loosely connected neuronal assemblies shows that they synchronize their sleep-like and awake-like states. Sleep probably evolved from a metabolically quiescent rest state. Any possibility of existence of a rest like state between sleep and awake states? Sleep seems to function to stabilize instinctual and learned memories. Sleep is a fundamental property of neuronal assemblies [16]. A network model for activity-dependent sleep regulation [17]. Organism sleep emerges from the local sleep states of functional units known as cortical columns; these local sleep states evolve through local activity inputs, loose couplings with neighboring cortical columns, and global regulation. Integration time scale is an important factor for state transition decisions. A minimum continuous sleep is necessary for memory consolidation [18]. Synapses in specific circuit become weaker and stronger during sleep [19]. Sleep might be regulated at a local level in the brain [20]. The role of sleep is to renormalize the synaptic weights after learning, resulting in a gain of synaptic strength in many brain circuits. Sleep can weaken most synapses but afford protection to some, including those directly activated by learning. How does selective synaptic down-selection occurs during sleep is not understood? It is unclear how and why some synapses are more selectively regulated than the rest [21].

Sleep regulatory mechanisms are neuronal in nature as seen traditionally. An alternative view is that glial cells, particularly astrocytes, may play a significant role in sleep regulation by forming feedback loops and globally regulating neuronal activity. [22]. Astrocytes dynamically change their activity across the sleep-wake cycle and may encode sleep needs via changes in intracellular signaling pathways. Astroglial sleep mechanisms are evolutionarily conserved. Astrocytes are dispersed widely in sub-cortical and cortical brain areas [23], including regions that trigger sleep and wakefulness. SCN astrocytes play a role in the circadian rhythm process [24].

Astrocytes modulate the accumulation of sleep pressure through a pathway involving A1 receptors. Astrocyte modulates behavior and provides strong evidence of the important role of A1 receptors in regulating sleep homeostasis and the cognitive decline associated with sleep loss. [25]. Astrocytes are involved in memory consolidation [26]. Astrocytic networks modulate neuronal activity, and behavior [27]. Long or short distance messaging is provided by the nonlinear gap junction in astrocytic network [28]. Astrocytic coverage of synapses increases during wake and reduces during sleep [29]. Astrocytes regulate cortical state switching playing a role in sleep, and regulation [30]. Astrocytes also promote sleep through the release of adenosine [31].

Despite many understandings so far from modeling perspective, several dynamics are missing and need to be incorporated in the two state model, such as continuous interaction of processes S and C, the interaction of SCN activity and sleep pressure, changes in the sleep pressure and its implications in clinical research [32]. Most recent outlook of the two process model, its importance, and implications were given in [33].

Taking inspiration from the hypothesized neuronal and astroglial feedback proposed model [34] for sleep homeostasis and considering it as a base for modeling. Model consists of three main components: First, neuronal waking signals that accumulate during the waking state reflect sleep needs. Second, an astroglial integrator of the neuronal waking signals. Third, astroglial derived sleep regulatory substances (SRS) that reduce the waking signals to promote sleep. Astrocytes may also form a key component of the sleep homeostat by integrating waking signals and releasing somnogenic substances and may explain both local and global aspects of sleep [34].

In this paper, author investigate the sleep awake dynamics initially with (i) a simple two ODE based homeostat model similar to two process model describing discrete state transitions in a continuous manner. Then, (ii) an astrocytic sleep pressure is introduced as a regulator giving rise to bistability and oscillatory behavior. Next, (iii) synaptic strength is modelled as a dependent function of sleep awake states. Lastly, (iv) stochastic version is considered to closely model the biological behavior.

## Methods

Python is used for writing the ODE and solving them numerically. XPPAUT used for plotting nullclines and bifurcation plots. Awake-sleep states are maintained by different awake and sleep-promoting neurons, respectively. Astrocytic sleep pressure regulates the concentration of sleep regulatory substances (SRS) [35].

## Results

### Sleep Awake Homeostat Model

Interaction between the two processes S and C regulate sleep awake dynamics. Processes S and C in the classical two process model interact with each other in a discrete manner [2]. Existing evidence indicates a continuous interaction exists between the circadian amplitude and homeostatic sleep pressure. High sleep pressure causes a lower circadian amplitude and vice versa. Also, SCN activity interacts with the sleep pressure. Circadian pacemaker obtains feedback from sleep homeostat such that increased sleep pressure reduces circadian amplitude. Strength of the clock output as a function of sleep homeostatic pressure. Thus, continuous interaction between the homeostatic and circadian process through a simple system of equations consisting of the two sleep and awake states is shown below:

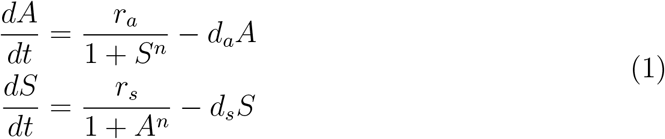

In the above system of equations 1, A/S represents the awake/sleep state maintained by the awake/sleep promoting neurons, *r_a_*/*r_s_* is the rate of entering into awake/sleep state, *d_a_/d_s_* is the rate of leaving from awake/sleep state, and n represents the nonlinearity exist in the system due to sleep/awake state promoting neuronal populations respectively. Sleep and awake state promoting neurons mutually inhibits each other in a way that the system 1 behaves as a toggle switch. It can dynamically switch between the discrete sleep-awake state similar to the flip-flop model [5] rather in a continuous manner. In figure (1A.), awake is a function of sleep. In figure (1B.), A is high and S is low. In figure (1C.) S is high, A is low. Figure 1 indicates state transitions occurs due to variations in the *d_a_* and *d_s_* state leaving rates in an alternative manner.

**Figure 1:**
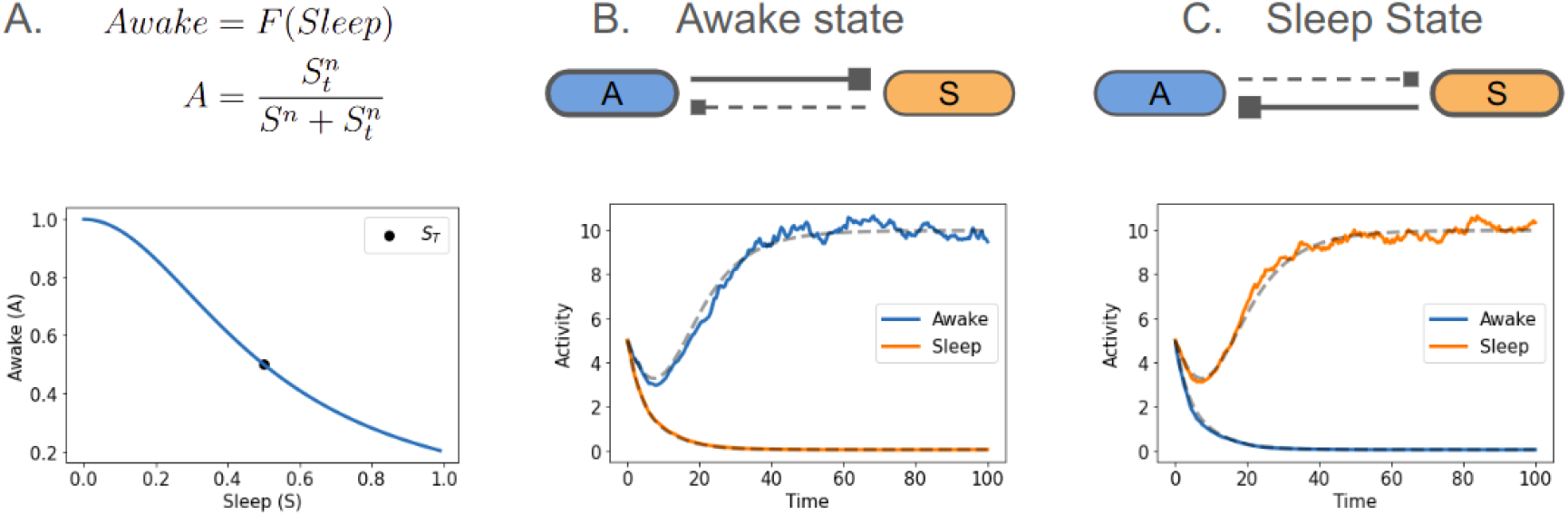
Sleep awake homeostat model circuit. (A.) Awake (*A*) state is a function of sleep (*S*) with sleep threshold (*S_t_*) where n represents the neuronal nonlinearity such that S inhibits A or vice versa. (B.) Awake state exists when *d_a_* < *d_s_* or low sleep pressure. (C.) Sleep state exists when *d_s_* < *d_a_* or high sleep pressure. Parameters for the model are: *S_t_*=0.5, n = 2 in (A.), *r_a_* = 1.0, *r_s_* = 1.0, n=2, (*d_a_* = 0.1, *d_s_*=0.2) in (B.), and (*d_a_* = 0.2, *d_s_*=0.1) in (C.)

The synaptic homeostasis hypothesis integrates sleep regulation with plasticity mechanisms [36]. Sleep is the price brain has to pays for synaptic plasticity. [37] Gain in the synaptic strength occurs during the awake state and loss during the sleep state, and despite selective synaptic strength are maintained [21] Glial cells in the fly brain also shows sleep homeostat dynamics [38]. Another proposed sleep homeostasis circuit encoding sleep pressure in Drosophila [39]. Hence, in the next section we introduce astrocytic sleep pressure regulating state transitions to the equations in 1.

### Sleep Awake Homeostasis with Sleep Pressure

Two state system mentioned in the system of equations 1 explains about the emergence of state transitions through time series simulations and phase plane analysis in a quantitative manner. But, it does not provide any insight into how these transitions occur in an autonomous manner and what processes govern the dynamics of such transitions as explained in the two process model? [2] Hence, an astrocytic sleep pressure (S) regulator shown in the figure 2 is added to the equations 1 as present below.

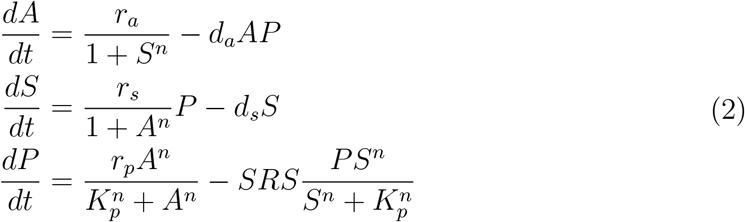

In system of equations 2, P represents the sleep pressure, *r_p_* is the rate of sleep pressure accumulation during awake state, *SRS* is the rate of sleep pressure degradation by sleep regulatory substances during sleep state, and *K_p_* is half of the pressure threshold. Sleep pressure (P) functions as a sleep integrator which negatively regulates the awake state during sleep accumulation and positively regulates the sleep state during sleep degradation.

**Figure 2:**
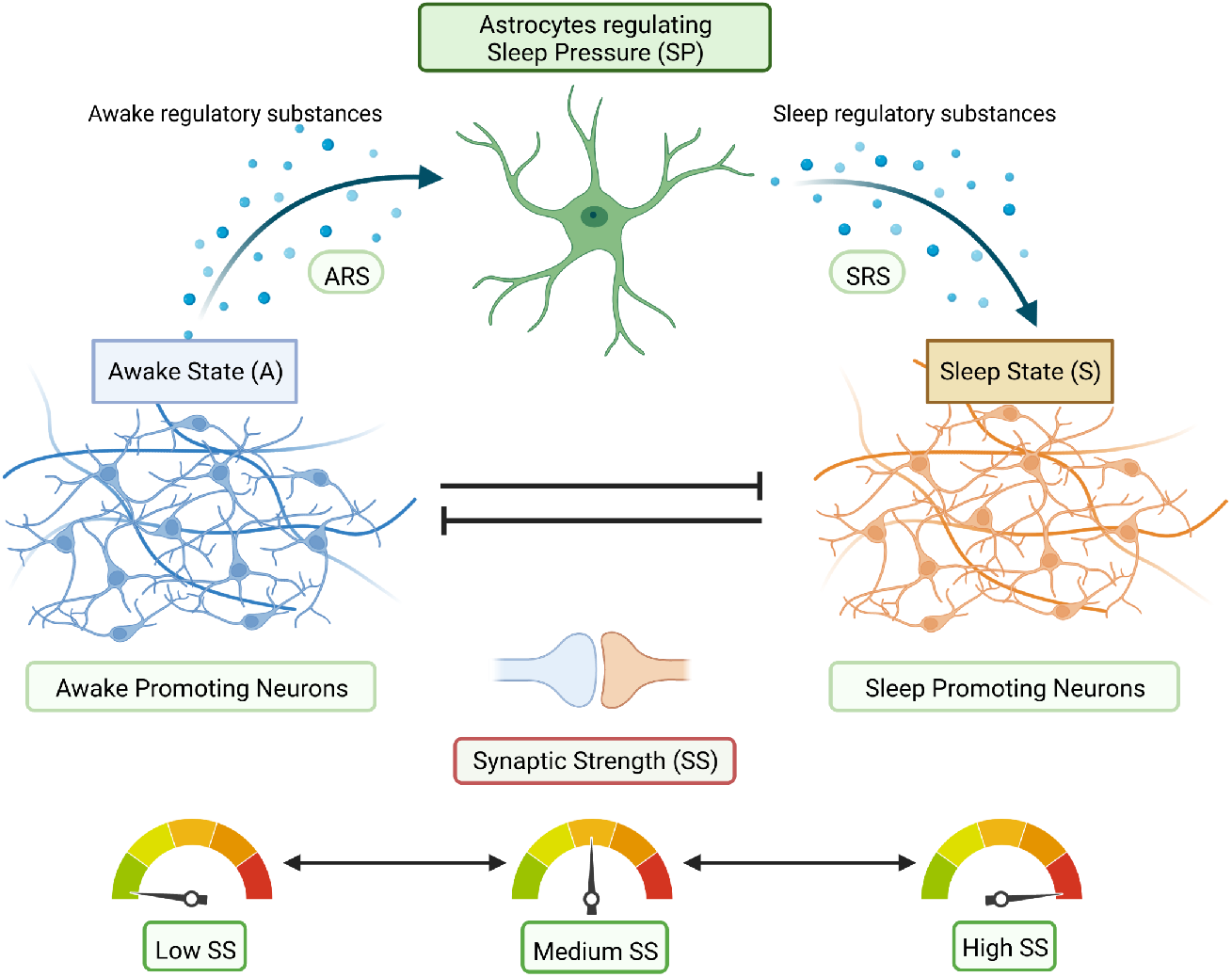
Astrocytic sleep awake homeostasis diagram. Created with BioRender.com A/S state promoting neurons mutually inhibits each other. During A state, both SP and SS are low. A inhibits S. SS increases and SP accumulates over the time causes release of SRS and beyond a threshold triggers state transition to S state. Meanwhile, both SP and SS are high. S inhibits A. SS decreases and SP degrades over time and with ARS below a threshold triggers a state transition from S to A.

Behavior of the system of equations 2 is shown in the figure 3. Figures (3A, 3B, and 3C) represent the circuit diagrams with feedback loops for normal, awake, and sleep state respectively. Time series data by varying initial conditions in figure (3D) shows presence of two stable states possibly representing sleep and awake states and an unstable state which might exist during sleep disorders. During awake state in (3B), incoherent feed-forward loop like circuit formation may be taking place by which A inhibits S and SP increases over the time. Phase plane analysis indicate bistability between the awake and sleep states with negligible pressure as shown in the Figure (3E) representing process S maintaining sleep homeostasis. During sleep state (3B), SP increases beyond a certain threshold may lead to a circuit transition from feedforward loop (FFL) to a negative feedback loop (FBL) which can produces oscillations in a continuous manner similar to process C maintaining circadian cycle.

**Figure 3:**
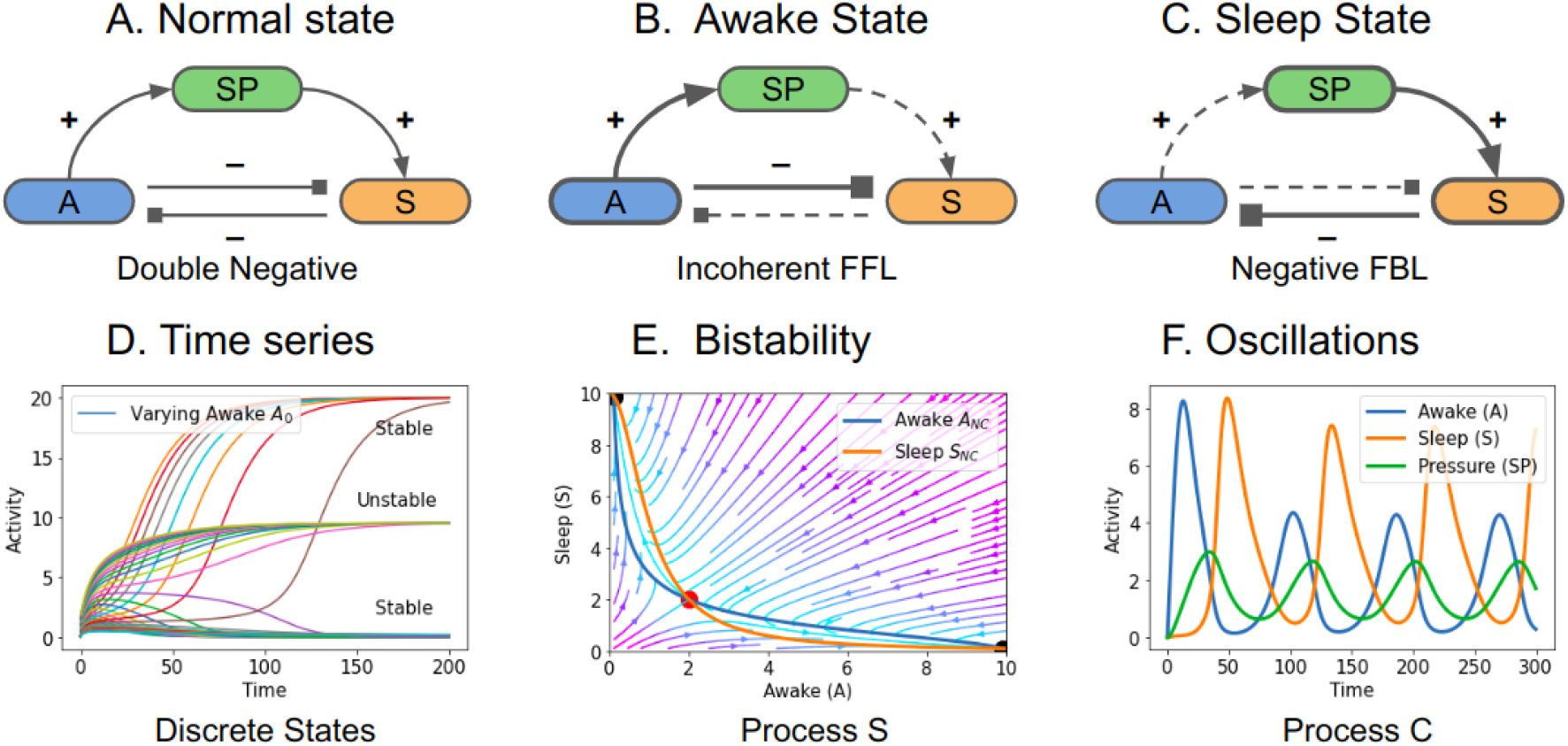
Sleep-awake states with sleep pressure dynamics. (A.) Mutually inhibiting sleep-awake states along with sleep pressure (SP) feedback. Initially, SP strength is negligible stem functions as a double negative circuit. (B.) A state inhibits the S state, and SP build-up takes place. Low SP during A state, system functions as an incoherent feedforward loop maintaining its state due to bistable nature of the circuit. (C.) High SP beyond the threshold results in a transition to S inhibiting A, and this negative feedback loop makes the circuit oscillate and reaches a stable homeostasis S state. (D.) Time series data indicating different states by keeping initial conditions (*S*_0_=0.1) and varying (*A*_0_) (E.) Bistability of the two-state model. (F.) Oscillatory behavior of the system. Parameter of the model: n=2, *r_a_*=1.0, *r_s_*=1.0, *d_a_*=0.1, *d_s_*=0.2, *r_p_*=0.2, *SRS*=0.1, *K_p_*=5.0 with initial condition (A=S=P=0).

### Sleep Awake Homeostasis along with Synaptic Strength

Synaptic strength varies globally during the awake and sleep cycle. It increases during the awake state and decreases during the sleep state. By further extending the model to understand how plasticity mechanisms are governed by synaptic strength modifications which occurs during sleep that can be both local or global [20]. A model of homeostatic regulation of connection strength and firing activity during sleep was proposed by [40]. The interplay between activity and plasticity during sleep can be understood by formulating a linear (Nonlinear) system in which the rate of change of synaptic strength (*S_s_*) depends linearly on neuronal activity *f* as firing rates. Plasticity mechanisms determine the decay rate of *S_s_* proportional to *f* = *aS_s_* where a is activity mechanisms and given by *dS_s_*/*dt* = –*Pf*. Therefore, we add synaptic strength in the system of equations 2 as a simple function of awake (A) and sleep (S) given by *S_s_*=*F*(*A, S*), and it is independent of the sleep pressure. Evolved system of equations to understand how synaptic strength is regulated during the state transition by astrocytic sleep pressure is mentioned below in 3.

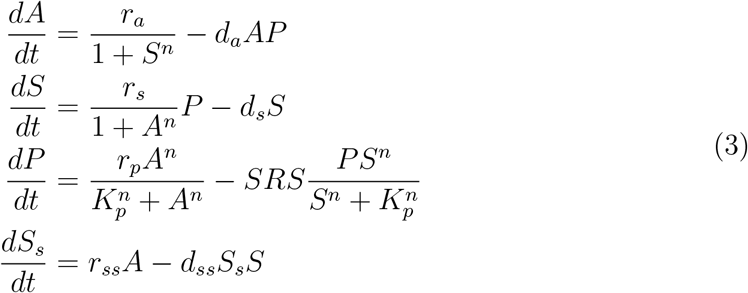

In equation 3, *S_s_* is the synaptic strength, *r_ss_* is the growth rate of synaptic strength during awake (A) state, and *dss* is the decay rate of synaptic strength during the sleep (S) state. Behavior of system of equations in 3 is given in the figure 4a. It is oscillatory in nature as seen earlier in previous figure 3 along with synaptic strength high during the awake state and low in sleep state. Bifurcation diagram is plotted between SP and SRS in the figure 4c. Sufficient release of SRS regulating the SP strength results in Hopf bifurcation which explains the emergence of a self oscillatory dynamics that exists between the sleep and awake state maintained by a lower (SRS>=0.06) and a upper (SRS<=0.3) threshold values. Simulations results matches with the experimental data [2] and indicates the importance of sleep pressure (SP) in sleep awake homeostasis and possibly as an independent astrocytic sleep pressure unit to maintain the synaptic strength locally or globally.

**Figure 4:**
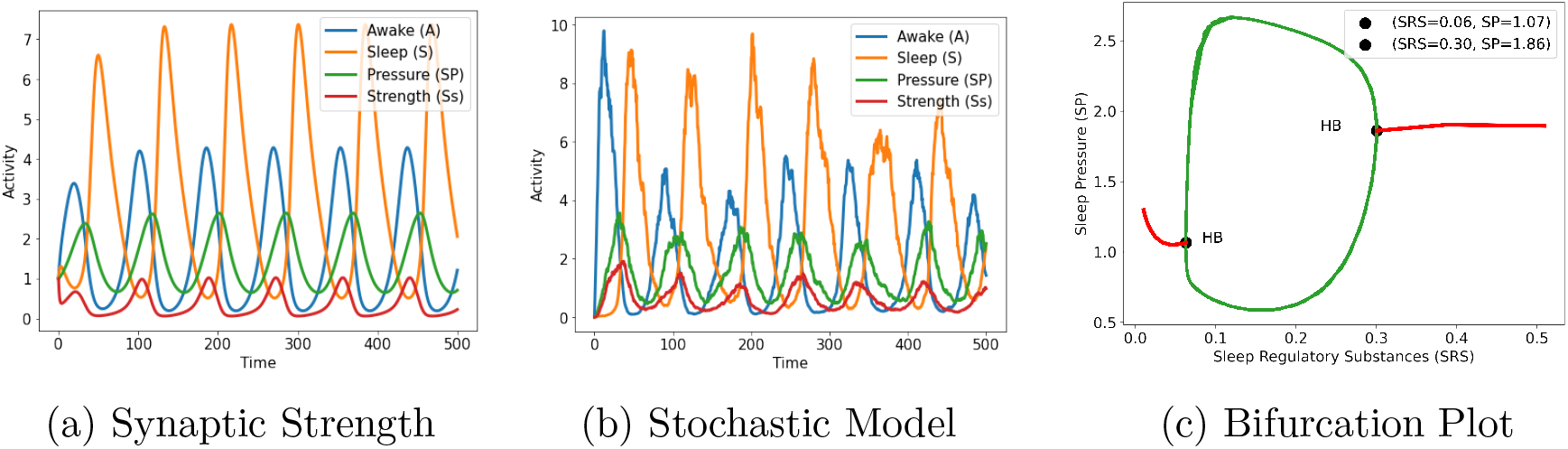
Sleep-Awake Homeostasis with Sleep Pressure and Synaptic Strength. (a). Model with synaptic strength. (b) Stochastic simulations of the astrocytic S/A model. (c). Hopf bifurcation plot between SP and SRS. SP regulates the sleep awake transition states by modulating SRS between two critical low and high threshold values. Parameters are same as given in the figure 3 with *r_ss_*=0.02 and *d_ss_*=1.1.

### Stochastic model of astrocytic sleep awake homeostasis

Sleep and awake state transitions are highly stochastic in nature due to the inherent noise present in the neuronal population level firing rates. Hence to incorporate the dynamics, a stochastic version of the previous deterministic model in equation 3 is simulated and its results are present in the figure 4b. Simulations results matches with the noisy firing patterns observed during the state transitions. Firing rates also increases or decrease during the transition [4]. Also, comparing the results with the experimental data [41], model explains how the firing rates are increased or decreased during the continuous state transitions as seen in the 4b with varying state amplitudes.

## Conclusions

In this research work, proposed sleep awake homeostasis model explained the dynamics of two state model [2] based on threshold and showed a relation between the process S and C in physiological manner. Explained the conditions required for dynamic transition between mutually inhibiting neuronal populations by time varying interactions for dynamic state switching through variations in SRS regulated by an astrocytic sleep pressure. Model explained state switching seen in the flip flop model [4]. Also captured the cycling behavior of reciprocal interaction model [1].

Sleep dysfunction or disorders may occur when system is not operating within the lower and upper sleep regulatory substances (SRS) thresholds. Model discussed about the existence of an unstable intermediate state which might possibly be present in the sleep disorders. Hence, the model can be used further to develop interventions for sleep disorders by modifying the SRS thresholds, sleep pressure strengths, or state promoting neuronal population activity for a specific sleep disorder. To study age-related changes in the sleep state using the model. Later it can be combined with the aging of the local circuits across the brain and how the elimination of dysfunctional reactive astrocytic sleep pressure units could result in maintaining cognitive health.

Model also explained about the global variations seen in the synaptic strength during sleep and awake states [37]. Astrocytes are involved in learning and memory consolidation during sleep. Astrocytes may selectively modify specific synapses to strengthen memory traces during memory consolidation. Also, plays a key role in general population level modification of synapses [37] to maintain a stable neuronal system. Astrocytes can implement selective plasticity for memory storage and general plasticity for functional needs by maintaining a distribution of neuronal firing rates [42]. Astrocytic sleep pressure may function as a global or local network of functional units connected through gap junctions which engage or disengage from the neuronal population in an independent manner and playing an important role in selective up-down regulation of synaptic strength [21] during awake or sleep states respectively.

Model integrates the sleep awake cycle with synaptic strength dynamics role in the consciousness research field. Proposed role of the SCN astrocytes in the circadian rhythm process [24] connects well with the missing piece of an astrocytic sleep pressure unit in the two process model. Hence, the study requires further investigations and experimental validations to support the hypothesis of an astrocytic sleep pressure functional unit in the sleep awake homeostasis across various model organisms.

Sleep is a fundamental property of neuronal assemblies such as cortical columns [16]. The fundamental units that transition between sleep and wake states (as reflected by functional changes in these units) are groups of tightly connected neurons in the cortex known as cortical columns. In the future, research work will focus upon understanding the principles of transition of a network of interacting neurons into a network of interacting assemblies [43] enabled by astrocytes. And develop a mathematical model by considering the sleep awake promoting neurons as neuronal ensembles regulated by an astrocytic sleep pressure network.

## Acknowledgement

This work is a part of the iCurious organisation to promote curiosity driven contribution to neuroscience. Standing on the shoulders of the pioneers, I thank all the researchers work that has made contribution in this article for conceptualization, modeling, simulation, and writing of the manuscript.

